# TIM-1 Serves as a Nonredundant Receptor for Ebola Virus, Enhancing Viremia and Pathogenesis

**DOI:** 10.1101/466102

**Authors:** Bethany Brunton, Kai Rogers, Elisabeth K. Phillips, Rachel B. Brouillette, Ruayda Bouls, Noah S. Butler, Wendy Maury

**Affiliations:** Department of Microbiology and Immunology, University of Iowa, Iowa City, IA 52242

## Abstract

**Background.:** T cell immunoglobulin mucin domain-1 (TIM-1) is a phosphatidylserine (PS) receptor, mediating filovirus entry into cells through interactions with PS on virions. TIM-1 expression has been implicated in Ebola virus (EBOV) pathogenesis; however, it remains unclear whether this is due to TIM-1 serving as a filovirus receptor in vivo or, as others have suggested, TIM-1 induces a cytokine storm elicited by T cell/virion interactions. Here, we use a BSL2 model virus that expresses EBOV glycoprotein and demonstrate the importance of TIM-1 as a virus receptor late during in vivo infection.

**Methodology/Principal findings.:** We used an infectious, recombinant vesicular stomatitis virus expressing EBOV glycoprotein (EBOV GP/rVSV) to assess the role of TIM-1 during in vivo infection. TIM-1-sufficient or TIM-1-deficient BALB/c interferon α/β receptor^-/-^ mice were challenged with EBOV GP/rVSV-GFP or G/rVSV-GFP. While G/rVSV caused profound morbidity and mortality in both mouse strains, TIM-1-deficient mice had significantly better survival than TIM-1-expressing mice following EBOV GP/rVSV challenge. EBOV GP/rVSV load in spleen was high and unaffected by expression of TIM-1. However, infectious virus in serum, liver, kidney and adrenal gland was reduced late in infection in the TIM-1-deficient mice, suggesting that virus entry via this receptor contributes to virus load. Consistent with higher virus loads, proinflammatory chemokines trended higher in organs from infected TIM-1-sufficient mice compared to the TIM-1-deficient mice, but proinflammatory cytokines were more modestly affected. To assess the role of T cells in EBOV GP/rVSV pathogenesis, T cells were depleted in TIM-1-sufficient and -deficient mice and the mice were challenged with virus. Depletion of T cells did not alter the pathogenic consequences of virus infection.

**Conclusions.:** Our studies provide evidence that at late times during EBOV GP/rVSV infection, TIM-1 increased virus load and associated mortality, consistent with an important role of this receptor in virus entry. This work suggests that inhibitors which block TIM-1/virus interaction may serve as effective antivirals, reducing virus load at late times during EBOV infection.

**Author summary:** T cell immunoglobulin mucin domain-1 (TIM-1) is one of a number of phosphatidylserine (PS) receptors that mediate clearance of apoptotic bodies by binding PS on the surface of dead or dying cells. Enveloped viruses mimic apoptotic bodies by exposing PS on the outer leaflet of the viral membrane. While TIM-1 has been shown to serve as an adherence factor/receptor for filoviruses in tissue culture, limited studies have investigated the role of TIM-1 as a receptor in vivo. Here, we sought to determine if TIM-1 was critical for Ebola virus glycoprotein-mediated infection using a BSL2 model virus. We demonstrate that loss of TIM-1 expression results in decreased virus load late during infection and significantly reduced virus-elicited mortality. These findings provide evidence that TIM-1 serves as an important receptor for Ebola virus in vivo. Blocking TIM-1/EBOV interactions may be effective antiviral strategy to reduce viral load and pathogenicity at late times of EBOV infection.

## Introduction

*Zaire ebolavirus* (EBOV) is one of five species of ebolaviruses within the *Filoviridae* family. EBOV continues to cause significant outbreaks in sub-Saharan Africa with case fatality rates as high as 90% [1]. All filoviruses have a broad species and cellular tropism. With the exception of lymphocytes, most cells within the body are thought to support EBOV infection and replication [2,3]. Histopathological studies of EBOV infected humans and non-human primates (NHPs) have demonstrated viral antigen in many different organs including: liver, spleen, lymph nodes, kidney, adrenal glands, lungs, gastrointestinal tract, skin, brain and heart [3-7].

A number of cell surface receptors are appreciated to mediate filovirus binding and internalization into the endosomal compartment of cells, including phosphatidylserine (PS) receptors [8,9] and C-type lectin receptors [10-14]. PS receptors do not interact with the viral glycoprotein (GP), but bind to PS on the surface of the virion lipid membrane, causing internalization of viral particles into the endosomal compartment [9,15]. This mechanism of viral entry has been termed apoptotic mimicry [16]. Following endosomal uptake of filovirions, proteolytic GP processing occurs, thereby allowing GP to interact with its endosomal cognate receptor, Niemann Pick C1 [17-21].

One important family of PS receptors is the T-cell immunoglobulin mucin domain (TIM) family. TIM family members, encoded by the *Havcr* family of genes, contribute to the uptake of apoptotic bodies to clear dying cells from tissues and the circulation [22-24]. TIM proteins are type 1, cell surface glycoproteins. Three family members are present in humans (hTIM-1, hTIM-3 and hTIM-4) and four in mice (TIM-1, TIM-2, TIM-3 and TIM-4) [25]. hTIM-1 was identified through a bioinformatics-based screen to be important for filovirus entry [8]. Subsequent studies demonstrated that hTIM-1 and hTIM-4, but not hTIM-3, enhance entry of a broad range of viruses including members of the alphavirus, arenavirus, baculovirus, filovirus, and flavivirus families [9,15,26-29]. Murine TIM-1 and TIM-4 also enhance enveloped virus uptake into the endosomal compartment [9,27,29].

The molecular interactions between TIM family members and enveloped viruses are well defined. The amino terminal IgV domain binds to PS on the outer leaflet of the viral membrane through a IgV domain binding pocket that is conserved across the TIM family of receptors [9,26,27,29]. Aspartic acid and asparagine residues within the binding pocket are essential for virion binding [9,15,27]; these same TIM residues are required for apoptotic body binding and uptake [30]. The IgV domain is extended from the plasma membrane by a mucin like domain (MLD) that is anchored to the cell surface with by a transmembrane domain connected to a short intracellular cytoplasmic tail. The length, but not the specific sequence, of the MLD is required for TIMs to serve as enveloped virus receptors [29]. Surprisingly, neither the TIM transmembrane domain nor cytoplasmic tail is required as a GPI-anchored TIM-1 construct is completely functional as a viral receptor [26,29]. These findings indicate that the TIM-1 cytoplasmic tail, which contains a tyrosine phosphorylation site that initiates signaling events [31-33], is not essential for TIM-1-mediated virus uptake.

While it is well established that TIM proteins serve as cell surface receptors for a number of enveloped viruses during in vitro infection of cultured cells, the importance of these family members for in vivo filovirus infection and pathogenesis has not been extensively examined. With the wide variety of cell surface receptors able to mediate filovirus uptake into endosomes, it is possible that sufficient receptor redundancy exists in vivo, such that the loss of any one of the PS receptors may have little or no effect on EBOV viremia, tissue virus load or pathological consequence. Alternatively, PS receptors are also immunomodulatory and been implicated in promoting inflammation. Thus, TIM proteins may exacerbate proinflammatory responses during virus infection. A recent study demonstrated that TIM-1-deficient mice have lower morbidity and mortality than wild-type mice when challenged intravascularly (i.v.) with mouse-adapted EBOV (maEBOV) [34]. This study highlighted the role of TIM-1 in non-permissive T lymphocytes, reporting that EBOV interaction with TIM-1 on CD4+ T cells enhanced proinflammatory cytokine dysregulation. The authors conclude that an enhanced TIM-1-dependent cytokine storm in T cells significantly contributes to EBOV pathogenesis. However, the impact of TIM-1 on virus load in mice was only examined in the plasma at a single time point, leaving open the possibility that TIM-1 may also serve as an important receptor for EBOV entry in vivo.

Here, we examined the in vivo importance of TIM-1 for virus replication and pathogenesis using a highly tractable BSL2 model virus of EBOV, which consists of recombinant vesicular stomatitis virus (VSV) encoding EBOV glycoprotein in place of the native VSV G protein (EBOV GP/rVSV). Our use of the EBOV GP/rVSV model virus allowed us to conduct detailed studies focused on the role of TIM-1 virus entry, host responses, and pathogenesis. As reported for maEBOV we observed that EBOV GP/rVSV was less pathogenic in TIM-1-deficient mice compared to control mice. The impact of the loss of TIM-1 was specific for EBOV GP-expressing virus since wild-type VSV was equally virulent in TIM-1-deficient and TIM-1-sufficient mice over a wide range of challenge doses. Importantly, reduced mortality observed in the virus-infected TIM-1^-/-^ mice was associated with lower virus load at late time points during infection in multiple tissues previously appreciated to be important in EBOV pathogenesis. Consistent with reduced overall virus loads, proinflammatory chemokine profiles were also lower in the EBOV GP/rVSV infected TIM-1-deficient mice at late time points following infection. Finally, to directly evaluate whether enhanced survival and reduced inflammation in TIM-1-deficient mice was associated with T cell activation as previously reported [34], we depleted T cells in EBOV GP/rVSV infected TIM-1-sufficient or -deficient mice and found that T cell depletion did not alter EBOV GP/rVSV pathogenesis. In total, our studies provide evidence that TIM-1 associated pathogenesis correlated with enhanced virus load at late times during infection, consistent with TIM-1 having an important role as a receptor for EBOV in vivo.

## Materials and Methods

### Ethics statement

This study was conducted in strict accordance with the Animal Welfare Act and the recommendations in the Guide for the Care and Use of Laboratory Animals of the National Institutes of Health (University of Iowa (UI) Institutional Assurance Number: #A3021-01). All animal procedures were approved by the UI Institutional Animal Care and Use Committee (IACUC) which oversees the administration of the IACUC protocols and the study was performed in accordance with the IACUC guidelines (Protocol #8011280, Filovirus glycoprotein/cellular protein interactions).

### Mice

BALB/c TIM-1-deficient mice have been previously described [35] and were a kind gift from Dr. Paul Rothman (Johns Hopkins University). Briefly, exons 4 and 5 of the TIM-1 gene, *Havcr1*, were replaced with a LacZ gene, generating a TIM-1-null mouse (TIM-1^-/-^) BALB/c IFN-αβ receptor-deficient (*Ifnar*^-/-^) mice were a kind gift from Dr. Joan Durbin, NYU Langone Medical Center. Mice were bred at the University of Iowa.

BALB/c *Ifnar*^-/-^ and BALB/c *Havcr1*^-/-^ (TIM-1^-/-^) mice were crossed for the creation of heterozygous progeny. Progeny were interbred and mice screened for the correct BALB/c *Ifnar*^-/-^/*Havcr1*^-/-^ genotype (referred to as TIM-1^-/-^ throughout this study). All expected genotypes were produced in normal Mendelian ratios. Genomic DNA from mouse tail-clips was assessed by PCR for genotypes. The primers and protocol for *Ifnar*^-/-^ screening has been previously described (218). *Havcr1* primer sequences included: shared forward, 5′ GTTTGCTGCCTTATTTGTGTCTGG 3′; WT reverse, 5′ CAGACATCA-ACTCTACAAGGTCCAAGAC 3′; knockout reverse, 5′ GTCTGTCCTAGCTTCCTCACTG 3′. PCR amplification was performed for 30 cycles at 94°C for 30 sec, 60°C for 30 sec, and 72°C for 1 min.

### Production of full length EBOV GP/rVSV virus and EBOV GPΔO/rVSV which lacked the mucin-like domain

These studies used recombinant, replication-competent vesicular stomatitis virus (VSV) expressing GFP and either full length EBOV GP (EBOV GP/rVSV-GFP) [18] (kind gift of Dr. Kartik Chandran), EBOV GP lacking the mucin domain of GP1 (EBOV GPΔO/rVSV-GFP)[8,15] or rVSV-GFP encoding its native glycoprotein, G (G/rVSV) (kind gift of Dr. Sean Whelan). Virus stocks were produced by infecting Vero cells, an African green monkey kidney epithelial cell line, at a low multiplicity of infection (MOI) of ~0.001 and collecting supernatants 48 hours following infection. Virus stocks were concentrated by centrifugation at 7,000 x g at 4°C overnight. The virus pellet was resuspended and centrifuged through a 20% sucrose cushion by ultracentrifugation at 26,000 rpm for 2 hours at 4°C in a Beckman Coulter SW32Ti rotor. The pellet was resuspended in PBS, treated with endotoxin removal agent (ThermoScientific #20339), aliquoted, and frozen at −80°C until use.

### Mouse infections

Five- to eight-week-old female BALB/c *Ifnar*^-/-^ (control) and BALB/c *Ifnar/Havcr1*^-/-^ (TIM-1^-/-^) mice were infected i.v. with recombinant, infectious VSV that encoded GFP and EBOV GP, EBOV ΔO or the native VSV G glycoprotein (EBOV GP/rVSV-GFP, EBOV GPΔO-rVSV-GFP and G/rVSV-GFP, respectively) using concentrations of virus noted in the figure legends. The dose of EBOV GP/rVSV or EBOV GPΔO-rVSV-GFP administered was dependent upon the stock. The dose of each stock was titered in vivo to give predictable high levels of mortality of control mice in 5-7 days. For studies with G/rVSV-GFP, either 10^1^ or 10^5^ iu of VSV virus was administered by i.v. injection. Survival was tracked; mice were weighed and scored for sickness daily. Clinical assessment of sickness was scored as follows: 0, no apparent illness; 1, slightly ruffled fur; 2, ruffled fur, active; 3, ruffled fur, inactive; 4, ruffled fur, inactive, hunched posture; 5, moribund or dead. While clinical assessments are not shown in figures, mice were humanely euthanized if they reached a score of 4. All mouse infection studies were concluded at 10 or 12 days following infection due to surviving mice regaining any lost weight and having no signs of clinical illness.

### Organ viral titers

Organs were harvested from control and TIM-1^-/-^ mice at 1, 3 or 5 days following infection from with EBOV GPΔO-rVSV. Prior to euthanasia, mice were anesthetized with isoflurane to perform retro-orbital bleeds for serum. Mice were euthanized and perfused with 10 mL of PBS through the heart and organs harvested, weighed and frozen at −80°C. To determine virus titers, organs or sera were thawed and organs were homogenized in PBS and filtered through a 0.45 μ syringe filter. Viral titers were determined by end point dilution on Vero cell as previously described [8]. Infection was scored 5 days following infection for GFP positivity using an inverted fluorescent microscope. Virus titers were calculated as 50% tissue culture infective dose (TCID50)/mL by the Spearman-Karber method. All organ titers were normalized according to the weight of the organ at harvest.

### Organ RNA isolation and reverse transcriptase quantitative PCR

Quantitative reverse transcriptase polymerase chain reaction (qRT-PCR) was used to detect proinflammatory cytokine and chemokines levels from organs of mice challenged with EBOV GPΔO/rVSV. At time of harvest, organs were placed in Trizol and frozen at −80°C until further use. Total RNA was isolated using TRIzol LS reagent (Life Technologies) according to manufacturer’s tissue RNA isolation procedure. RNA was quantified by Nanodrop (Thermo Scientific). Total RNA (2 μg) was reverse transcribed into cDNA using random primers and the High-Capacity cDNA Reverse Transcription kit (Applied Biosystems). SYBR Green based quantitative PCR reactions (Applied Biosystems) were performed using 1.5μL of a 1:100 dilution of cDNA from each reaction and specific primers for murine cytokines and chemokines. Primer sequences are found in Supplemental Table I. Expression levels of the cytokine/ chemokines of interest were defined as a ratio between threshold cycle (Ct) values for the gene of interest and the endogenous control, mouse HPRT, and is displayed as the log2 value of this ratio.

### T cell depletion studies

Five- to eight-week-old female BALB/c *Ifnar*^-/-^ and BALB/c *Ifnar*/TIM-l^-/-^ mice were injected with 200μg of anti-CD4 (clone GK1.5) and 200μg anti-CD8 (clone 2.43) depleting monoclonal antibodies both one day prior to retroorbital infection with EBOV GP/rVSV-GFP and two days post infection. Survival was tracked; mice were weighed and scored for sickness daily as described above to assess euthanasia criteria for each infected mouse. Prior to infection with EBOV GP/rVSV-GFP, depletion was validated by isolating peripheral blood mononuclear cells from both depleted and non-depleted animals and staining of PBMCs with anti-CD90 antibody (clone 30-H12). Staining was done by incubating with anti-CD90 antibody in FACS buffer and Fc block (2.4G2) for 30 minutes, washing 3 times to remove excess antibody, and detecting fluorescence on a BD FACSCalibur.

### Statistics

Statistical analyses were performed using GraphPad Prism software (GraphPad Software, Inc.). Results are shown as means or geometric means and standard error of the means (s.e.m.) or geometric s.e.m., respectively, is shown where appropriate. Log-rank (Mantel-Cox) tests were used to analyze differences in survival. In vivo experiments were performed at least in duplicate with at least 8 mice total per treatment group. Mice or samples were randomly assigned to various treatment groups. All data points and animals were reported in results and statistical analyses. For the nonparametric viral titer data, Mann-Whitney U-test was used. *P* values less than 0.05 were considered significant. For two way comparisons between control and experimental values, a Student’s t-test was performed.

## Results

### TIM-1 expression enhances EBOV GP/rVSV infection, but not VSV

To create a TIM-1 deficient mouse, exons 4 and 5 of the *Havcr1* gene encoding TIM-1 were replaced with the LacZ gene by homologous recombination as previously described [35]. This mouse strain was used to study the role of TIM-1 in allergic airway diseases and Th2 responses [35]. Phenotypic characterization of TIM-1^-/-^ mice revealed no differences in immune cell numbers, immune system development, or immunological homeostasis compared to WT mice [35]. BALB/c TIM-1^-/-^ mice were bred onto a BALB/c interferon αβ receptor (*Ifnar*^-/-^) knock out background since type I interferon abrogates replication of the BSL2 recombinant EBOV GP/rVSV used in these studies [36,37]. Homozygous BALB/c *Ifnar*/TIM-1^-/-^ and *Ifnar*^-/-^ mice (called TIM-1^-/-^ and control mice, respectively, throughout the remainder of this study) were used for all infections. Challenge virus was administered intravenously since this route of delivery mimics a primary route of EBOV transmission, blood-to-blood contact. Control and TIM-1^-/-^ mice were challenged with the lowest dose of virus that produced predictable death in control mice in 5-7 days (Supplemental Fig. 1). Minor titer variations were observed between virus stocks and dosages were adjusted accordingly.

We challenged the TIM-1^-/-^ and control mice with full length EBOV GP/rVSV or EBOV GPΔO/rVSV, which has the GP1 mucin like domain (MLD) deleted. EBOV GPΔO pseudovirions and recombinant viruses have the same tropism as virus bearing EBOV GP [8,38-40]. Use of both viruses in these studies allowed us to determine if the elimination of the mucin domain altered the pathogenesis associated with in vivo challenge with these viruses. As expected, TIM-1-sufficient control mice succumbed to EBOV GP/rVSV or EBOV GPΔO/rVSV between days 4-7 of infection (Fig. 1A and B). By contrast, TIM-1^-/-^ mice challenged with the same dose had significantly reduced mortality following EBOV GP/rVSV or EBOV GPΔO/rVSV infection and delayed time-to-death of those that did succumb to infection. Weight loss associated with infection in the TIM-1^-/-^ mice were also significantly reduced at some days (data not shown). These findings indicate that TIM-1^-/-^ mice had improved survival when infected with EBOV GP/rVSV compared to controls and that survival was not affected by the presence of the GP1 MLD.

**Fig. 1.**
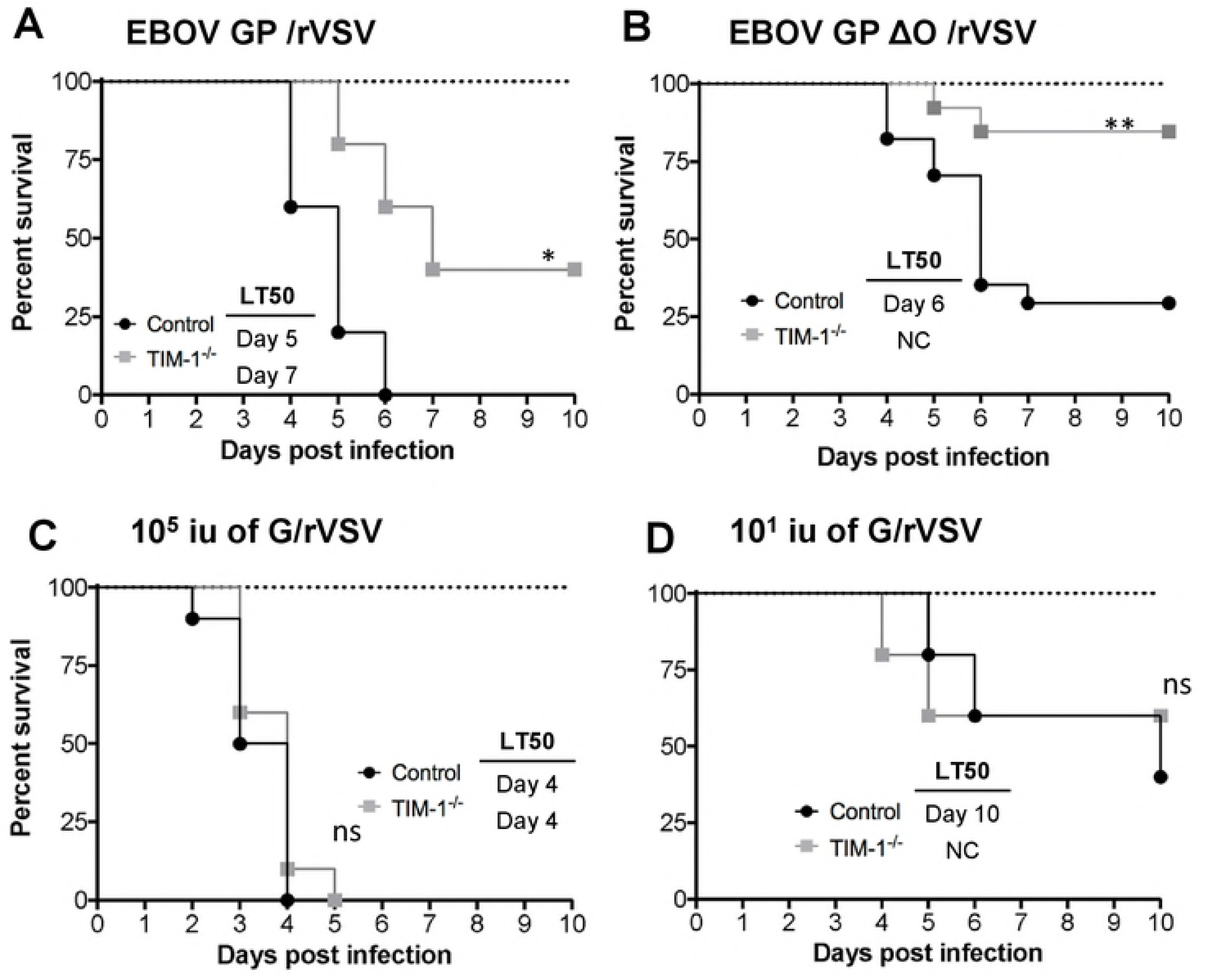
Loss of TIM-1 reduces mortality following EBOV GP/rVSV and EBOV GP ΔO/rVSV infection, but not G/rVSV. A and B. Female BALB/c *Ifnar*^-/-^ (control) and BALB/c *Ifnar*/TIM-1^-/-^ (TIM-1^-/-^) mice infected with 10^5^ iu EBOV GP/rVSV (A; n = 5 mice per group) or EBOV GP ΔO/rVSV (B; n = 13-17 mice per group) by intravenous injection. C. Female BALB/c *Ifnar*^-/-^ (control) and BALB/c *Ifnar*/TIM-1^-/-^ (TIM-1^-/-^) mice infected with 10^5^ iu G/rVSV (n = 10 mice per group). D. Similar G/rVSV challenge studies as shown in panel C, but mice were challenged with 10^1^ iu (n = 5 mice per group) of G/rVSV. Survival was assessed following infection for all mouse studies. Significance for survival curve was determined by Log Rank (Mantel-Cox) test, ^*^ p< 0.05, ^**^p < 0.01. LT50 = median lethal time until death; NC = noncalculable.

In tissue culture studies, we have shown that hTIM-1 does not mediate WT VSV entry [8], presumably because the cognate receptor for VSV, LDL receptor is abundantly present on target cells and mediates VSV entry [41]. However, the relevance of TIM-1 in vivo for VSV infection has not been examined. Further, WT VSV serves as an excellent control for in vivo studies with EBOV GP/rVSV. We challenged TIM-1^-/-^ and control mice with 10^5^ iu of VSV by i.v. injection.

In contrast to our EBOV GP/rVSV findings, we observed no difference in the survival curve between the two strains of mice (Fig. 1C). Since it is likely that VSV bearing it native GP is more pathogenic than a recombinant VSV containing a different viral GP, we also evaluated mortality associated with different doses of VSV and found that administration of as little as 10^1^ iu of VSV was lethal to *Ifnar*^-/-^ mice (Supplemental Fig. 2). Thus, we repeated VSV infections in control and TIM-1^-/-^ mice at a challenge dose of 10^1^ iu to determine if subtle changes in virus pathogenesis could be discerned. Even at this low dose, there was no difference in the survival in the TIM-1^-/-^ mice versus the control mice (Fig. 1D). These results provide evidence that the difference in EBOV GP/rVSV pathogenesis in BALB/c *Ifnar*^-/-^ and TIM-1^-/-^ mice was due to the presence of EBOV GP expressed in the recombinant VSV rather than other VSV genes. The reduced pathogenesis of EBOV GP expressing virus in TIM-1^-/-^ mice was consistent with findings described by Younan et al. using maEBOV [34].

### Murine TIM-1 enhances EBOV GP/rVSV virus load at late times during infection

The effect of TIM-1 expression on viremia and organ viral loads following i.v. EBOV GPΔO/rVSV infection was examined in serum and organs harvested 1, 3 or 5 days following infection (Fig. 2). Viremia and infectious virus in various organs were quantified by endpoint dilution titering on Vero cells, a highly permissive cell line for EBOV GP/rVSV. At early times during infection, no difference in viremia or virus load was observed in most organs of TIM-1 versus control mice. However, by day 5 of EBOV GPΔO/rVSV infection, TIM-1^-/-^ mice had a 100-fold reduction in viremia compared to control mice (Fig. 2) and a similar trend was observed during infections with full length EBOV GP/rVSV (Supplement Fig. 3). In parallel, levels of infectious virus in liver, kidney, and adrenal gland were also significantly reduced. Studies at day 5 of infection also indicated that EBOV GPΔO/rVSV loads were much reduced in the brain of TIM-1^-/-^ mice and trended lower in the testis (Supplement Fig. 4A and B), consistent with an overall reduction in virus load in the TIM-1^-/-^ mice at late times during infection. Thus, reduced virus replication in a number of organs was associated with the survival observed in TIM-1^-/-^ mice. These findings provide evidence that TIM-1 expression is important for the generation of high viral load in some organs at late times in infection.

**Fig. 2.**
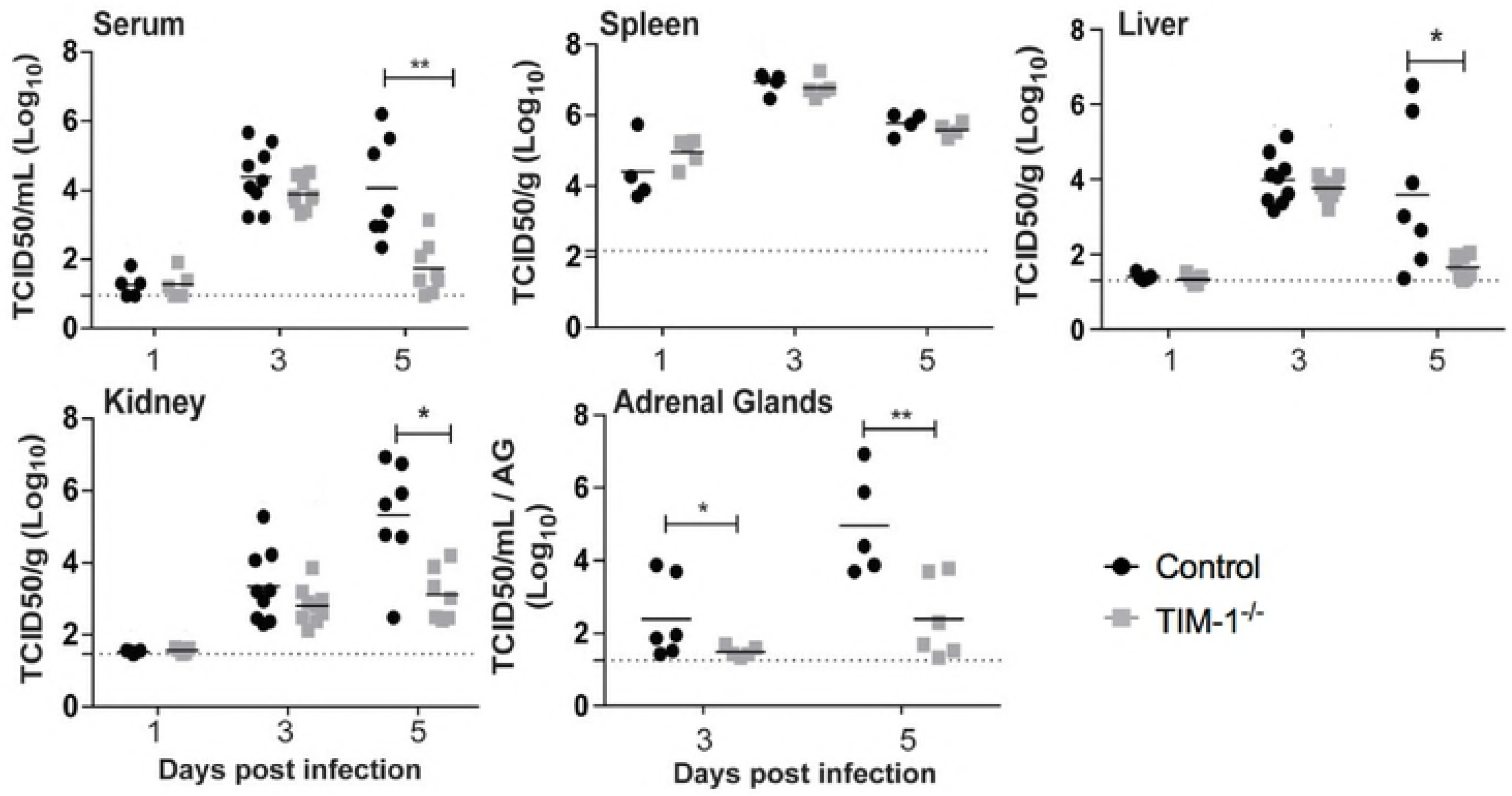
Reduced viremia and viral loads in a variety of organs of TIM-1^-/-^ mice at late time points following i.v. EBOV GP ΔO/rVSV infection. Sera and organs were harvested from BALB/c *Ifnar* ^-/-^ (control) and BALB/c *Ifnar*/TIM-1^-/-^ (TIM-1^-/-^) mice at days 1, 3 and 5 following infection with 10^5^ iu of EBOV GP ΔO /rVSV by i.v. injection. Titers were determined by endpoint dilution of serum or homogenized organ samples on Vero cells. Solid lines indicate geometric mean for each data set. Dotted line indicates the level of detection. Adrenal gland (AG) titers are displayed as per AG homogenized in 1 ml of PBS. Significance was calculated by Mann-Whitney test to compare control to TIM-1^-/-^ mice at each time point, ^*^*p* < 0.05, ^**^*p* < 0.01. ns, not significant.

Viral loads in the spleen and lungs were not affected by the loss of TIM-1 (Fig. 2 and Supplemental Fig. 4C). The viral burden in the spleen was significantly higher at day 1 than in any other organ assessed and remained high in both mouse strains throughout the course of infection with a peak in titers occurring at day 3. These results are consistent with previous studies that implicate spleen in early and sustained EBOV replication [42-44]. Lung titers were not significantly different between the control and TIM-1^-/-^ mice at 5 days following infection. This result was somewhat unexpected as we had previously demonstrated robust hTIM-1 expression on the apical surface airway epithelial cells [8]. As TIM-1 was not observed to be expressed on the basolateral side of lung epithelium, TIM-1 may be important for entry of aerosolized EBOV entry into a host, but may not influence basolateral infection of lung via the circulation.

### TIM-1-expressing mice exhibit elevated levels of specific proinflammatory chemokines following EBOV GP/rVSV infection

Elevated proinflammatory and immunomodulatory cytokines and chemokines are evident in serum and infected organs during EBOV infection of animal models and patients [45-51]. To determine if reduced virus load in TIM-1-deficient mice at late time points was associated with lower RNA expression profiles of selected, well-characterized cytokines, levels in the spleen, liver and kidney were examined prior to and following EBOV GPΔO/rVSV infection. Organs were harvested at day 3 and 5 following infection and total RNA was isolated and amplified for the mRNA of the housekeeping gene, HPRT, and the cytokines IL-6, TNF, IL-12 and IL-10. Cytokine expression levels were normalized against HPRT expression. Overall, baseline values of the organ cytokine expression from uninfected control and TIM-1^-/-^ mice were low with little difference between the strains (Fig. 3). While at day 5 TNF was significantly higher in spleen of control mice, in general during the infection cytokine expression was variable within groups and levels were not significantly different between the two strains of mice.

**Fig. 3.**
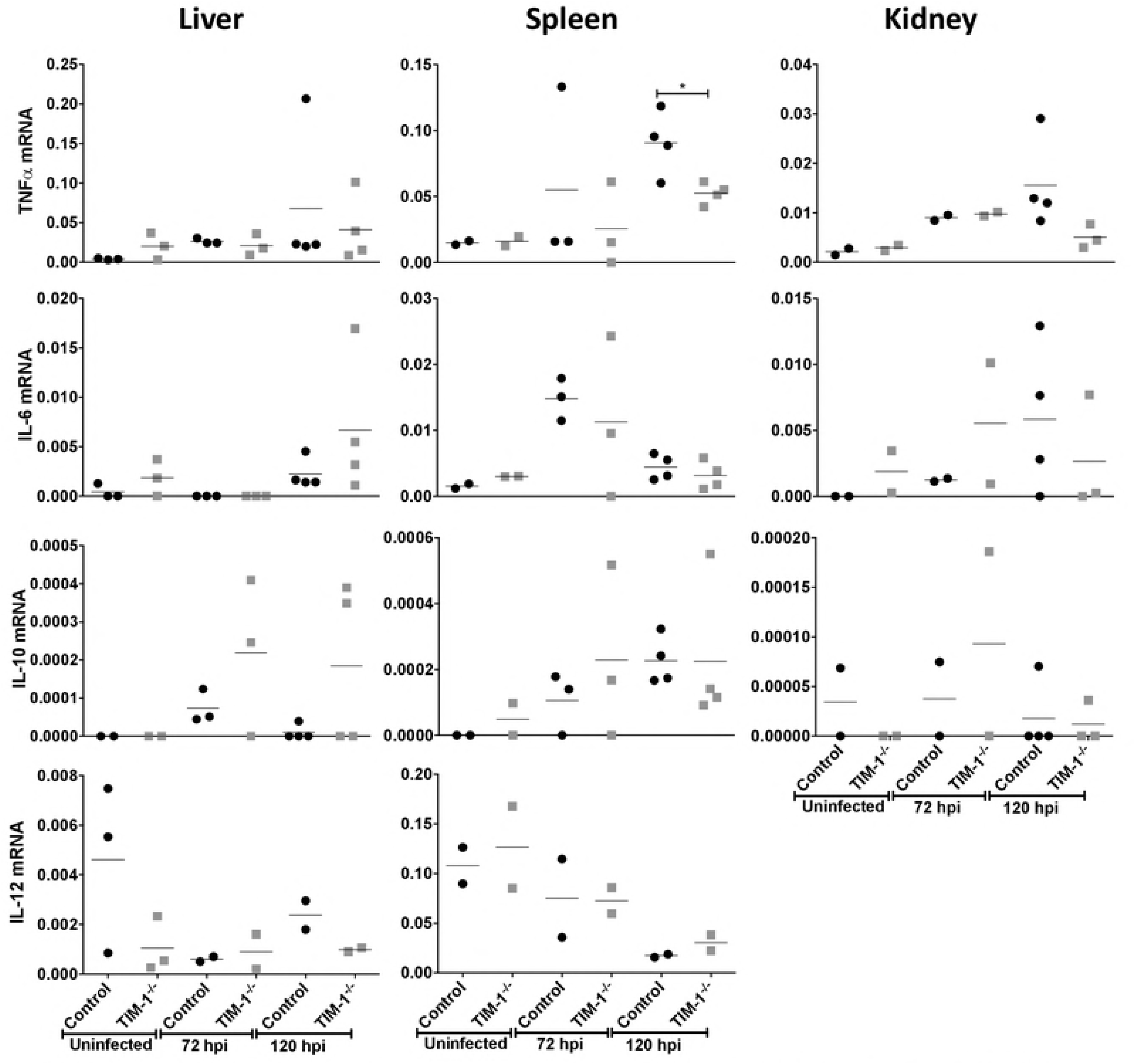
Cytokine expression in liver, spleen and kidney of EBOV GPΔO/rVSV-infected *Ifnar*^-/-^ and *Infar*/TIM-1^-/-^ mice. Tissues were harvested from uninfected and infected BALB/c *Ifnar* ^-/-^ (control) and BALB/c *Ifnar* /TIM-1^-/-^ (TIM-1^-/-^) mice. For infection studies, tissues were harvested at 3 or 5 days following infection with 10^5^ iu of EBOV GPΔO/rVSV by i.v. injection. RNA was isolated from the organs and expression of mouse TNF, IL-6, IL-10 and IL-12, were quantified by qRT-PCR. Results represent cytokines expression relative to murine HPRT for at least three independent livers, spleens and kidneys. Data points represent values for individual mice. Solid lines indicate the mean for each data set. Statistical significance was determined by Student’s t-test compared the control mice for each time point and is only shown for those comparisons observed to differ, ^*^p<0.05.

Elevated levels of several chemokines and growth factors have been implicated in fatal EBOV disease outcomes including MIP-1α, MIP-1β, MCP-1, M-CSF, MIF, IP-10, GRO-α and eotaxin [49]. Therefore, we analyzed control and TIM-1^-/-^ organs following EBOV GP^-/-^O/rVSV infection for the chemokines, CXCL10 (IP-10) and CCL2 (MCP-1). At least one of the two chemokine transcripts in all three organs was elevated in the control mice at both day 3 and/or 5 of infection compared to the TIM-1^-/-^ mouse tissues (Fig. 4). In combination with our survival and viral burden results, these observations suggest that the presence of TIM-1 in mice contributes to EBOV GP/rVSV pathogenesis through increased infection of cells in several organs at late times during infection and that this is associated with increased expression of proinflammatory chemokines.

**Fig. 4.**
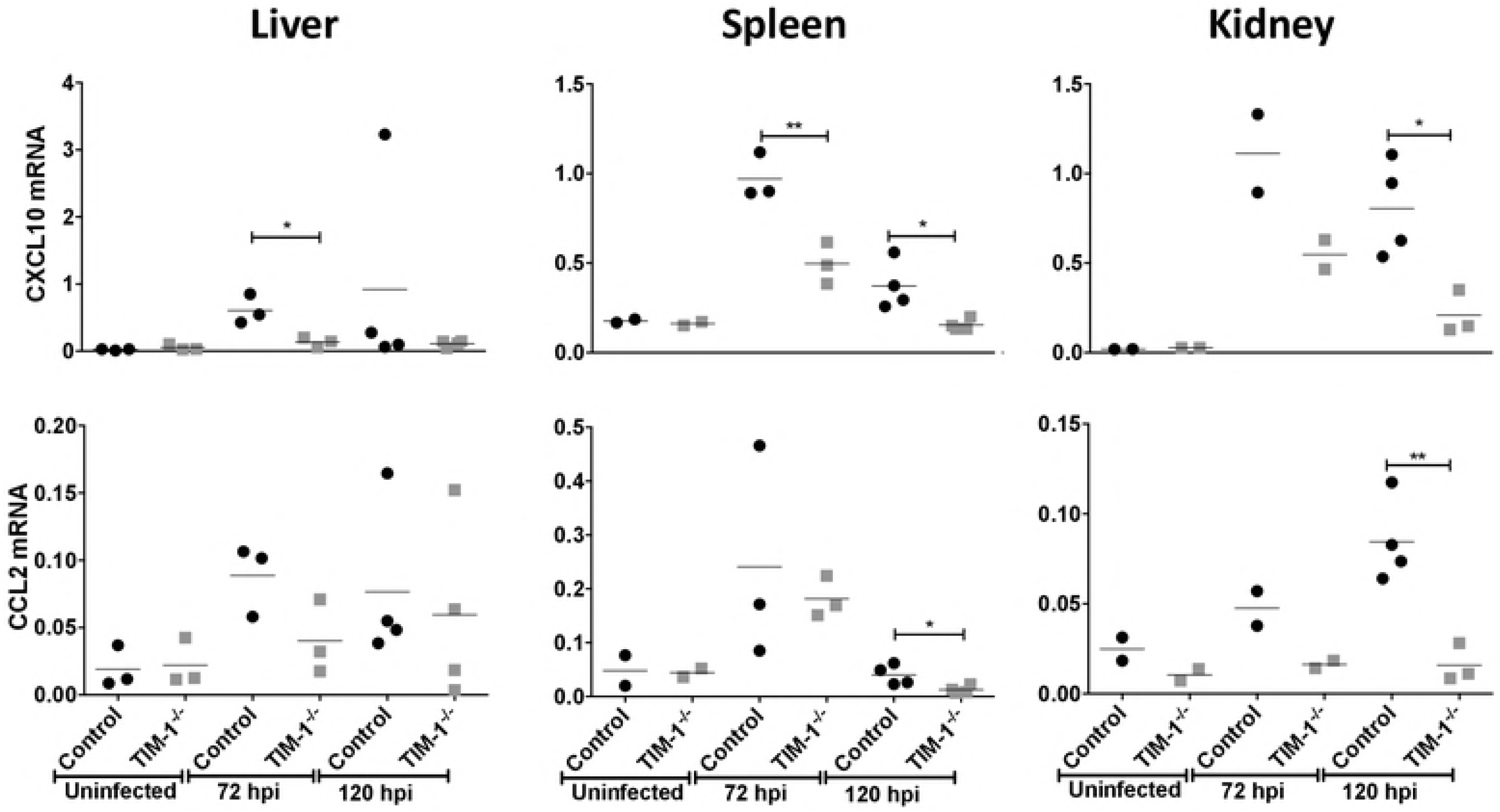
Chemokine CXCL10 and CCL2 expression in the liver, spleen and kidney of EBOV GPΔO/rVSV-infected control and TIM-1^-/-^ mice. Tissues were harvested from uninfected and infected BALB/c *Ifnar*^-/-^ (control) and BALB/c *Ifnar*/TTM-1^-/-^ (TIM-1^-/-^) mice. For infection studies, tissues were harvested at 3 or 5 days following infection with 10^5^ iu of EBOV GPΔO/rVSV by i.v. injection. RNA was isolated from the organs and proinflammatory chemokines, mouse CXCL10 and CCL2, were quantified by qRT-PCR. Results represent chemokine expression relative to murine HPRT for at least three independent livers, spleens and kidneys. Data points represent values for individual mice. Solid lines indicate the mean for each data set. Statistical significance was determined by Student’s t-test compared the control mice for each time point and is only shown for those comparisons observed to differ, ^*^p<0.05.

### T cell depletion does not alter morbidity associated with EBOV GP/rVSV infection

TIM-1 is expressed by a range of hematopoietic and non-hematopoietic cells [52]. This study showed that virus load in spleen, an organ rich in hematopoietic cells, was not affected by the loss of TIM-1 expression, suggesting that it might be TIM-1 expression on non-hematopoietic cells late during infection that affects EBOV GP/rVSV load and survival. As others have suggested that TIM-1 on T cell subsets may contribute to enhanced EBOV pathogenesis [34], we depleted T cells in control and TIM-1^-/-^ mice to assess outcomes during EBOV GP/rVSV infection. Mice were intraperitoneally administered α-CD8 mAb, 2.43, and α-CD4 mAb, GK1.5, at days −1 and 2. We verified that T cells within peripheral blood were profoundly depleted at day 5 of infection by flow cytometry following immunostaining with an α-CD90 mAb (Fig. 5A). As observed for the T cell-competent mice in above studies, T cell-depleted control mice challenged with EBOV GP/rVSV succumbed to the infection between 4-6 days. Likewise, while T cell-depleted TIM-1^-/-^ mice had significantly better survival than T cell depleted control mice they did not exhibit improved survival over TIM-1^-/-^ mice that were not T cell-depleted (Fig. 5B). These data provide evidence that the presence of T cells does not alter the course of this acute infection and suggests that TIM-1 expression on non-T cell populations contributes to pathogenesis.

**Fig. 5.**
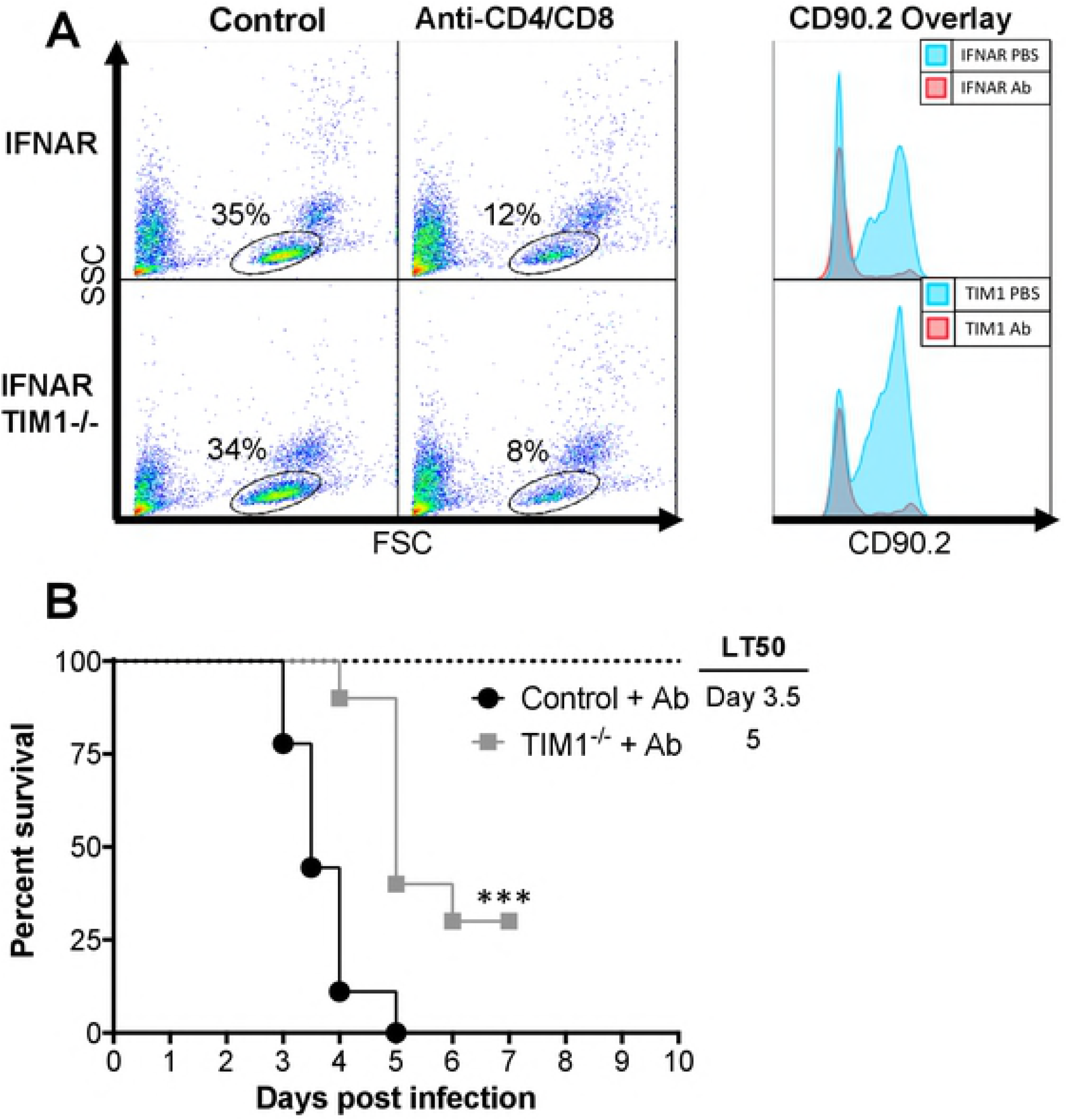
T cell depletion does not alter the survival protection conferred by the loss of TIM-1 expression. A. Intraperitoneal injection of α-CD8 mAb, clone 2.43, and α-CD4 mAb, clone GK1.5, treatment at days −1 and 2 systemically depletes T cell populations in Female BALB/c *Ifnar*^-/-^ (Control) and BALB/c *Ifnar*^-/-^ /TIM-1^-/-^ (TIM-1^-/-^) mice as determined by α-CD90 mAb staining of peripheral blood mononuclear cells at day 5 following EBOV GP/rVSV infection. CD90.2 overlay depicts the subset of cells gated in the panel on the left.
B. Survival was assessed following infection with 7×10^2^ iu of EBOV GP/rVSV administered by intravenous infection (n = 10 mice per group) and two treatments of α-CD8 mAb and α-CD4 mAb at Day −1 and 2 from infection. Significance for survival curve was determined by Log Rank (Mantel-Cox) test, ^***^p < 0.001.

## Discussion

Here, we show that loss of TIM-1 expression decreased overall mortality and delayed time-to-death of those mice that do succumb when challenged with EBOV GP/rVSV. The impact on survival of TIM-1 expression was similar with rVSV bearing MLD-deleted EBOV GP, indicating that the presence of the MLD did not affect the observed pathogenesis. Consistent with the enhanced survival of the TIM-1-deficient mice following virus challenge, we show that these mice also had reduced infectious virus in liver, kidney and adrenal gland at late times during infection. EBOV replication in these organs is well established and is thought to contribute to overall EBOV load [42, 53-55]. The lower virus load in these organs of the TIM-^-/-^ mice was also reflected in a ~100-fold reduction in serum viremia at day 5 of infection. The reduced pathology in our TIM-1-deficient mice was EBOV GP-dependent since survival associated with G/rVSV infection was unaffected by TIM-1 expression. Thus, our studies indicate that the glycoprotein present on the virions was responsible for the TIM-1-dependent changes in virus load and mouse survival.

Interestingly, we did not observe that all organs previously implicated as important in EBOV infection had lower virus load in TIM-1^-/-^ mice. Splenic viral loads were at high throughout infection in both control and TIM-1^-/-^ mice. These data suggested that TIM-1 expressing cells do not appreciably contribute to splenic virus loads and that splenic loads can be high in mice without those animals necessarily succumbing to infection.

While the TIM-1 does not interact directly with EBOV GP, the binding of TIM-1 to virion-associated PS has been shown to elicit viral particle entry into the endosomal compartment [9,15] where EBOV GP is proteolytically processed, binds to NPC1 and mediates membrane fusion [17,18,20,21,56-58]. Filoviral particle entry into endosomes occurs through interactions with a number of other cell surface receptors in tissue culture. However, these studies and those by Younan, et al [34] demonstrate that in vivo at least one receptor, TIM-1, is not redundant with other receptors, but uniquely contributes to EBOV pathogenesis. Future studies to evaluate the role of additional host receptors implicated in EBOV entry would provide valuable insights to the potential receptor redundancy. These receptors include other TIM family members, TAM tyrosine kinase receptors and C-type lectins.

The correlation between enhanced survival and reduced viral loads in the TIM-1^-/-^ mice suggests that TIM-1 serves as a virus receptor for EBOV in some organs late during infection. Likely, late cell targets that express TIM-1 would include kidney epithelial cells [59] and epithelial cells in other organs [8]. Surprisingly, decreased virus load was not observed in an earlier study that challenged TIM-1^-/-^ mice with maEBOV [34]. Instead, the authors reported that the genome copy number in plasma did not significantly differ in TIM-1-sufficient and - deficient mice. However, in this study, virus load was shown from a single time point during infection. The discrepancy between our findings and the previous study may be due to the tissues examined and/or the timing of the sampling.

The physiological role of TIM-1 has been extensively studied. Agonistic monoclonal antibody binding to TIM-1 on CD4^+^ T, iNKT and splenic B cells induces cellular activation in a wide range of organisms from zebrafish to humans [24,32,33,59-61]. This observation has led to the understanding that TIM-1 serves as a costimulatory molecule on these cells and leads to upregulation of cytokines in T and iNKT cells [24,59], as well as antibody production by B cells [61]. In contrast, transient TIM-1 expression on injured kidney epithelial cells serves an anti-inflammatory role through its uptake and clearance of apoptotic bodies [62].

Younan, et al. described the role of TIM-1 in EBOV pathology to TIM-1 stimulation of T cell cytokine and chemokine dysregulation [34]. Yet proinflammatory cytokines were only modestly altered in our studies even at late times during infection in TIM-1-sufficient mice. We did observe elevated levels of the proinflammatory chemokines, CCL2 and CXCL10, in the TIM-1-sufficient mice compared to the deficient mice and postulated that the higher levels of chemokines in TIM-1^+^ mice may reflect the innate immune responses stimulated by the higher virus load. Alternatively, as postulated by Younan, et al., the elevated chemokine profile and associated mortality in the TIM-1^+^ mice might be due to a TIM-1-dependent cytokine storm elicited by T cells [34]. We tested this latter possibility by virus challenge of T cell-depleted mice. T cell depletion did not alter EBOV GP/rVSV pathology. Significantly greater mortality was associated with virus infection of TIM-1-sufficient mice which were depleted for T cells than T cell-depleted, TIM-1-deficient mice, suggesting that T cells are not responsible for the reduced survival of TIM-1-sufficient mice. Hence, our findings do not support the conclusion that TIM-1 expression on T cells plays a significant role in the pathology associated with this acute infection.

Our studies and studies performed by Younan et al. [34] delivered EBOV GP/rVSV intravenously. In studies not shown, we observed that intraperitoneal (i.p.) delivery of EBOV GP/rVSV or maEBOV into WT versus TfM-1^-/-^ mice was equally pathogenic. This finding may be explained by the previous observation that another TIM family member, TIM-4, is highly expressed on resident peritoneal macrophages [63] and is used as a receptor for EBOV [27]. Likely, the use of TIM-4 as a receptor within this compartment usurps the need for TIM-1 expression during i.p. challenge, even late during infection.

Our results also demonstrate that TIM-1 is not important for WT VSV pathogenesis. Due to the wide cellular tropism of VSV, ubiquitous cell lipid components like PS, phosphatidylinositol or the ganglioside GM3 were originally proposed as the VSV cell surface receptor [64-66]. However, more recent investigations have revealed that these lipids are not readily used as VSV cell surface receptors [67,68]. Instead, the LDL receptor and its family members are proposed to serve as VSV receptors on human and mouse cells [41]. Therefore, in vivo pathogenesis induced by VSV would differ from EBOV GP/rVSV since the dependence on LDL receptors for entry is conferred by the VSVG glycoprotein [41]. Presumably the VSV membrane contains PS that can interact with TIM-1, but the affinity of VSV G for LDL receptors is likely greater than the affinity of PS in the virion envelope towards PS receptors like TIM-1. Studies from our lab have shown that only when the high affinity interactions of Lassa virus GP with its receptor, α-dystroglycan, are abrogated does TIM-1 mediate Lassa virus pseudovirion entry [69]. Future studies would be valuable to assess the ability of VSV to utilize TIM-1 as a cell surface receptor in the absence of expression of LDL receptors.

Liver and kidney dysfunction and necrosis are integral aspects of EBOV pathology of humans, NHPs [3,70] and mice [71,72]. Our studies indicate that TIM-1 expression is associated with elevated viral loads in the liver, kidney, adrenal gland, and brain since loss of TIM-1 significantly lowered viral burden in these organs. Future studies will need to explore the impact that TIM-1 expression has on EBOV infection of specific cells within these organs. By identifying TIM-1 expressing cells that serve as viral targets and understanding the contribution of these cells to the EBOV disease pathogenesis, we will be able to better develop TIM-1 specific therapeutics against EBOV infection.

## Acknowledgements

We would like to thank Drs. Al Klingelhutz and Patrick Sinn for their helpful comments on the manuscript.

## Supplemental data

**Supplemental Table 1. Cytokine/Chemokine primer sequences for qRT-PCR analysis. Supplemental Fig. 1.** Mortality (A, C) and weight loss (B, D) associated with increasing doses of EBOV GPΔO/rVSV (A,B) and EBOV GP/rVSV (C,D). All virus was administered iv. n=1-3 mice per group.

**Supplemental Fig.2. Weight loss following intraperitoneal infection of *Ifnar* ^-/-^ mice with 10-/fold serial dilutions of VSV.** BALB/c *Ifnar* ^-/-^ mice (1-4 mice per dose) received the indicated dose of EBOVΔO GP/rVSV virus by i.p. injection. Weight loss was tracked over 10-days to determine the lowest predictably lethal dose (10^1^ infectious units). Grey lines indicate the virus doses that caused mortality in all or some of the mice over the course of the experiment with 100% of mice succumbing to the 10^1^ iu dose.

**Supplemental Fig. 3. EBOV/rVSV serum titers.** Sera were harvested from BALB/c *Ifnar* ^-/-^ (control) or BALB/c *Ifnar*^-/-^ /TIM-1^-/-^ (TIM-1^-/-^) mice at days 1, 3 and 5 following infection with 10^5^ iu of EBOV GP/rVSV by i.v. injection. Titers were determined by endpoint dilution of serum on Vero cells. Solid lines indicate geometric mean for each data set. Dotted line indicates the level of detection. Significance was calculated by Student’s t-tests comparisons of the geometric means.

**Supplemental Fig. 4. Reduced viral loads in the brain but not lungs of *Ifnar*^-/-^ /TIM-1^-/-^ mice 5 days following i.v. EBOV GP ΔO /rVSV infection.** Brain (A), testis (B) and lung (C) tissue were harvested from BALB/c *Ifnar*^-/-^ (control) to BALB/c *Ifnar* ^-/-^ /TIM-1^-/-^ (TIM-1^-/-^) mice at day 5 following infection with 10^5^ iu of EBOV GP ΔO /rVSV by i.v. injection. Titers were determined by endpoint dilution of homogenized organ samples on Vero cells. Shown are data points for individual mice within each treatment and the bold line represent the mean titers from serum of of 2-4 mice per group.

